# Real-time projections of epidemic transmission and estimation of vaccination impact during an Ebola virus disease outbreak in Northeastern Democratic Republic of Congo

**DOI:** 10.1101/461285

**Authors:** Lee Worden, Rae Wannier, Nicole A. Hoff, Kamy Musene, Bernice Selo, Mathias Mossoko, Emile Okitolonda-Wemakoy, Jean Jacques Muyembe-Tamfum, George W. Rutherford, Thomas M. Lietman, Anne W. Rimoin, Travis C. Porco, J. Daniel Kelly

## Abstract

**Background:** As of February 25, 2019, 875 cases of Ebola virus disease (EVD) were reported in North Kivu and Ituri Provinces, Democratic Republic of Congo. Since the beginning of October, the outbreak has largely shifted into regions in which active armed conflict has occurred, and in which EVD cases and their contacts have been difficult for health workers to reach. We used available data on the current outbreak with case-count time series from prior outbreaks to project the short-term and long-term course of the outbreak.

**Methods:** For short- and long-term projections, we modeled Ebola virus transmission using a stochastic branching process that assumes gradually quenching transmission rates estimated from past EVD outbreaks, with outbreak trajectories conditioned on agreement with the course of the current outbreak, and with multiple levels of vaccination coverage. We used two regression models to estimate similar projection periods. Short- and long-term projections were estimated using negative binomial autoregression and Theil-Sen regression, respectively. We also used Gott’s rule to estimate a baseline minimum-information projection. We then constructed an ensemble of forecasts to be compared and recorded for future evaluation against final outcomes. From August 20, 2018 to February 25, 2019, short-term model projections were validated against known case counts.

**Results:** During validation of short-term projections, from one week to four weeks, we found models consistently scored higher on shorter-term forecasts. Based on case counts as of February 25, the stochastic model projected a median case count of 933 cases by February 18 (95% prediction interval: 872–1054) and 955 cases by March 4 (95% prediction interval: 874–1105), while the auto-regression model projects median case counts of 889 (95% prediction interval: 876–933) and 898 (95% prediction interval: 877–983) cases for those dates, respectively. Projected median final counts range from 953 to 1,749. Although the outbreak is already larger than all past Ebola outbreaks other than the 2013–2016 outbreak of over 26,000 cases, our models do not project that it is likely to grow to that scale. The stochastic model estimates that vaccination coverage in this outbreak is lower than reported in its trial setting in Sierra Leone.

**Conclusions:** Our projections are concentrated in a range up to about 300 cases beyond those already reported. While a catastrophic outbreak is not projected, it is not ruled out, and prevention and vigilance are warranted. Prospective validation of our models in real time allowed us to generate more accurate short-term forecasts, and this process may prove useful for future real-time short-term forecasting. We estimate that transmission rates are higher than would be seen under target levels of 62% coverage due to contact tracing and vaccination, and this model estimate may offer a surrogate indicator for the outbreak response challenges.

**Author summary:** As of February 25, 2019, 875 cases of Ebola virus disease (EVD) were reported in North Kivu and Ituri Provinces, Democratic Republic of Congo. Since the beginning of October 2018, the outbreak has largely shifted into regions in which active armed conflict has been reported, and in which EVD cases and their contacts have been difficult for health workers to reach. We used an ensemble of models to estimate EVD transmission rates and to forecast the short- and long-term course of the outbreak. Our models project that a final size of roughly up to 300 additional cases is most likely, and estimate that transmission rates are higher than would be seen under optimal levels of contact tracing and vaccination. While a catastrophic outbreak is not projected, is it not ruled out, and prevention and vigilance are warranted.

## Introduction

On August 1, 2018, the World Health Organization (WHO) announced a new outbreak of Ebola virus disease (EVD) in North Kivu Province, Democratic Republic of Congo (DRC) [1]. Epidemiological investigations traced EVD cases back to the week of April 30 and identified the initial epicenter to be Mabalako. This region of DRC (Fig 1) has over five million inhabitants, some of whom suffer from armed conflict, humanitarian crisis, and displacement to the bordering countries of Uganda and Rwanda [2]. Since the outbreak began, its magnitude has surpassed all other EVD outbreaks in DRC, becoming the second largest in history to date. As of February 25, 2019, 875 EVD cases had been reported (65 probable and 810 confirmed), and the Ministry of Health of DRC, the World Health Organization, and other organizations were responding to the Ebola outbreak [3].

**Fig 1.**
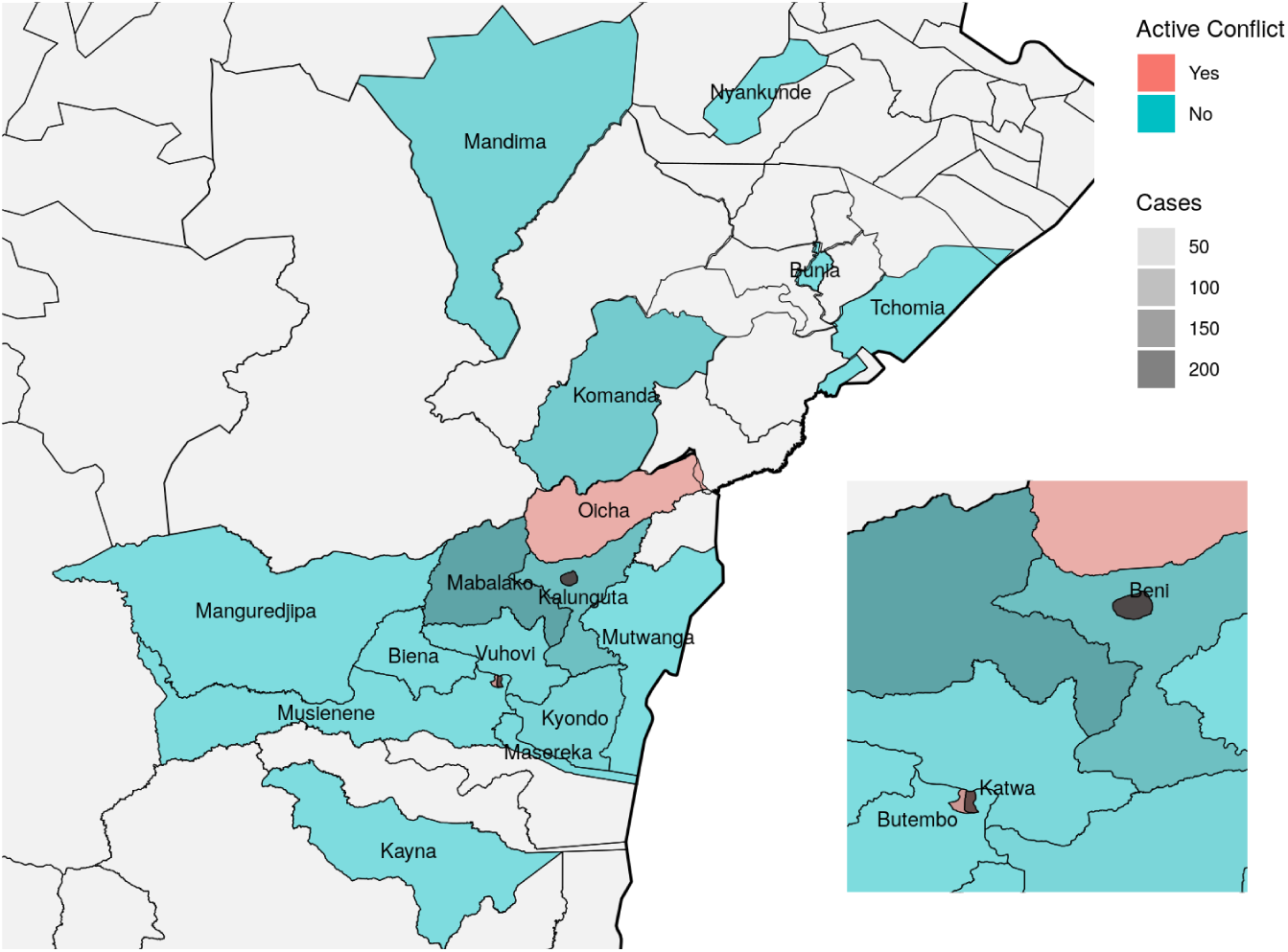
**Map of health districts in which confirmed and/or probable cases have occurred** in Northeastern DRC, with zones in which active conflict has occurred identified by color. Data sourced from Référentiel Géographique Commun [7], made available under the Open Data Commons Open Database License.

This is the first EVD outbreak to occur in an armed conflict zone, and this new context has created challenges to the outbreak response [4]. Uptake of traditional control measures such as contact tracing and safe burials has varied with the community, even with the rapid deployment and widespread availability of experimental vaccines and therapeutics [3]. Our understanding of the impact of scientific advances on EVD transmission has been evolving; there has been an underlying, untested assumption that vaccination may reduce epidemic transmission [5]. Mathematical modeling of simulated EVD epidemics suggested that vaccination coverage as low as 40% in the general population and 95% in healthcare workers (HCWs) may avert another epidemic similar to size of the West African one [6]. Studies have yet to estimate the levels of vaccination coverage associated with epidemic decline. An even greater potential problem in difficult-to-access outbreak areas is ascertainment of vaccination coverage among contacts and contacts of contacts. Mathematical models are needed that can produce relatively accurate forecasts in the setting of such unknown but important outbreak response metrics.

During an Ebola outbreak, real-time forecasting has the potential to support decision-making and allocation of resources, but highly accurate forecasts have proven difficult for Ebola [8, 9] as well as other diseases [10–13]. Highly accurate forecasts of small, noisy outbreaks may be a fundamentally elusive ideal [14]. Previous work has found that probabilistic forecasts can have relatively high accuracy within a few weeks, though they become less useful as time horizons grow longer [15]. Thus, short-term forecasts may provide useful information for response organizations.

In this paper, we applied a suite of independent methods to estimate short- and long-term projections of future EVD case counts in northeastern DRC in real time. Given the unique context of the current EVD outbreak, we validated short-term projections before generating projections on February 25. One of our models also estimated vaccination coverage. It is our hope that these projections offer insight into the outbreak trajectory and vaccination coverage, particularly in the setting of armed conflict.

## Methods

We used four independent methods to generate projections of future case counts: (1) a stochastic simulation model, (2) a negative binomial auto-regression model, (3) a Theil-Sen regression model, and (4) a final size projection based on Gott’s rule. The stochastic simulation model was calibrated to time-dependent transmission rates measured from past EVD outbreaks and constrained to the observed partial trajectory of the current outbreak, extending a model used in previous work [16]. This model generated short- and long-term projections. The negative binomial auto-regression model produced short-term projections based on case counts from the outbreak to date and prior outbreaks. The Theil-Sen regression model predicted final outbreak size based on prior outbreaks. We used a probability mass function derived from Gott’s rule as a minimum-information projection of final outbreak size, using only the assumption that the proportion of the outbreak observed so far is entirely unknown. From these, we assembled an ensemble of projections to be compared and recorded for future evaluation against final outcomes.

### Data sources

Data on the current outbreak were collected from the WHO website in real time as updated information was published [3]. A cumulative case count of probable and confirmed cases was extracted to be consistent with the best knowledge at the time. Snapshots of the table of case counts were kept as of multiple dates (Fig S1 in S1 Supporting Information), to be used in retrospective scoring of model projections against subsequently known counts. Although the epidemic was officially reported on August 1 as a cluster of cases occurring in June and July, seven sporadic early cases from April and May were subsequently linked to the current outbreak and were added to later case totals [3]. This additional knowledge was added retrospectively to the time series of cumulative case counts only for predictions made for days on or after September 15, when these cases were officially linked to the current outbreak.

### Stochastic model

We modeled Ebola virus (EBOV) transmission using a stochastic branching process model, parameterized by transmission rates estimated from the dynamics of prior EVD outbreaks and conditioned on agreement with reported case counts from the 2018 EVD outbreak to date. We incorporated high and low estimates of vaccination coverage into this model. We used this model to generate a set of probabilistic projections of the size of simulated outbreaks in the current setting. This model is similar to one described in previous work [16], with the addition of a smoothing step allowing for a continuum of transmission rates interpolated between those estimated from prior outbreaks.

On the assumption that past outbreaks provide a basis for projection of the current outbreak, we used estimates of transmission rates from past EVD outbreaks to parameterize simulations of the current outbreak. To estimate the reproduction number *R* in past outbreaks as a function of the number of days from the beginning of the outbreak, we included reported cases by date from fourteen prior outbreaks (Table S1 in S1 Supporting Information), [17–30]. To reflect the Ebola response system in DRC during what is now its tenth outbreak, the first historical outbreak reported in each country was excluded (e.g. the 1976 outbreak in Yambuko, DRC), as there is a difference in the Ebola response system as well as community sensitization to EVD following a country’s first outbreak. We used the Wallinga-Teunis technique to estimate *R* for each case and therefore for each reporting date in these outbreaks [31]. The serial interval distribution used for this estimation was a gamma distribution with a mean of days and a standard deviation of 5 days, with intervals rounded to the nearest whole number of days, consistent with the understanding that the serial interval of EVD cases ranges from 3 to 36 days with mean 14 to 15 days.

Transmission rates estimated by day in these outbreaks tend to decline from initially high to eventually low values, though they may display substantial fluctuations. This “quenching” of transmission may be driven by formal interventions such as quarantine, or by informal changes in individuals’ behavior in response to the disease or by depletion of uninfected contacts of infective individuals, or by other causes. We incorporated this pattern into our model by estimating an initial reproduction number *R*_initial_ and quenching rate *τ* for each outbreak by fitting an exponentially quenched curve to the outbreak’s estimates of *R* by day *d* (Fig S2 in S1 Supporting Information), and used these pairs of parameters, one from each past outbreak, to construct a joint distribution of initial reproduction numbers and quenching rates for outbreak simulation.

We simulated EBOV transmission using a stochastic branching process model in which the number of secondary cases caused by any given primary case is drawn from a negative binomial distribution, whose mean is the reproduction number *R* as a function of day of the outbreak, and variance is controlled by a dispersion parameter *k* [32, 33]. All transmission events were assumed to be independent. The interval between date of detection of each primary case and that of each of its secondary cases is assumed gamma distributed with mean 14.5 days and standard deviation 5 days, rounded to the nearest whole number of days, as above.

We used the (*R*_initial_, *τ*) pairs estimated from prior outbreaks to provide *R* values for simulation. *R*_initial_ values were sampled uniformly from the range of values estimated from past outbreaks. We applied a linear regression to the values of *R*_initial_ and log(*τ*) estimated for prior outbreaks and used the resulting regression line to assign a mean *τ* to each *R*, used with the residual variance of log(*τ*) as a distribution from which to sample *τ* values for simulation given *R*_initial_. Note that the range of fast and slow quenching scenarios modeled in this way is not limited to the exact combinations estimated from past outbreaks, but extends over a continuous distribution that includes those values. The parameters *R*_initial_ and *τ* sampled in this way, together with three values of the dispersion parameter *k*, 0.3, 0.5, and 0.7, consistent with transmission heterogeneity observed in past Ebola outbreaks, were used to generate simulated outbreaks.

This model generated randomly varying simulated outbreaks with a range of case counts per day. The outbreak was assumed to begin with a single case. The simulation was run multiple times, each instance producing a proposed epidemic trajectory, generated by the above branching process with the given parameters *R*_initial_, *τ*, and *k*, and these were then filtered by discarding all proposed trajectories except those whose cumulative case counts matched known counts of the current 2018 EVD outbreak on known dates. In earlier, smaller data sets we filtered against all reported case counts, while in later, more complete data sets we used a thinned series of case counts for filtering, for computational tractability, by selecting five case counts evenly spaced in the data set plus the final case count (Fig S1 in S1 Supporting Information). The filtration required an exact match of the first target value, and at subsequent target dates accepted epidemics within a number of cases more or less than each recorded value. On the earlier data sets in which the beginning dates of the epidemic were unknown, the first target value was allowed to match on any day, and subsequent target dates were assigned relative to that day.

In order to produce model outbreak trajectories consistent with the case counts reported more recently in the outbreak, it was necessary to make the filtering step of the model more tolerant to variation in counts in order to accommodate the rapidly rising count. This was because higher transmission rates in late September and early October were necessary to generate case counts of that size than were consistent with the earlier counts.

Thus this model embodies a set of assumptions that transmission rates are overall gradually declining from the start of the outbreak to its end, though possibly with a high level of variability in transmission rate between cases and between simulations. When the tolerance of the filter on case counts is small, quenching of transmission through time must closely track case counts, while when tolerance is high, fluctuations in the rate of generation of new cases can reflect a pattern of ongoing quenching of transmission more loosely and on the long term, while being more insensitive to short-term up and down fluctuations in transmission rates reflected in the true case counts.

We varied the tolerance as the data set became more complete to maintain a roughly fixed rate of generation of filtered trajectories. As larger tolerances became necessary, in data sets from after October, we introduced one further distinction: while it is possible for cumulative case counts to decrease as inaccurately classified cases are removed from the counts, due to the precision of the labeling of cases probable and confirmed we expect this to happen rarely, so we limit the tolerance of matching to only at most 15 cases below the reported count regardless of the tolerance of counts above the reported count. This limit on underestimates was applied only to analysis of data sets from later than October 13, to preserve unaltered the projection methods we reported in a preprint of this paper [34]. Filtering tolerances were as follows: when using the August 20 data set, we allowed a tolerance of 4 cases more or less than each target count; for August 27 and September 5, 6 cases; for September 15, 10 cases; for October 7, 12 cases; for October 13, 17 cases; for November 1, 41 cases; for November 20, 55 cases; for January 6, 2019, 75 cases; and for February 25, 150 cases. This one-step particle filtering technique produced an ensemble of model outbreaks, filtered on agreement with the recorded trajectory of the outbreak to date. This filtered ensemble was then used to generate projections of the outbreak in the short term (one week, two weeks, four weeks, and eight weeks) and long term (final size) [35].

To model vaccination coverage with respect to total transmission (unreported and reported), we multiplied the estimate of vaccine effectiveness by low and high estimates of reported cases. In a ring vaccination study at the end of the West Africa outbreak, the overall estimated recombinant vesicular stomatitis virus (rVSV)-vectored vaccine efficacy was 100% and vaccine effectiveness was 64.6% in protecting all contacts and contacts of contacts from EVD in the randomized clusters, including unvaccinated cluster members [36]. We used estimates of vaccine effectiveness in our stochastic model. The ring vaccination study found the vaccine to be effective against cases with onset dates 10 days or more from the date of vaccine administration, so we modeled the vaccination program as a proportionate reduction in the number of new cases with onsets 10 days or more after the program start date.

We used past estimates of the proportion of unreported cases to estimate the proportion of exposed individuals not covered by the vaccination process. Based on a Sierra Leonean study from the 2013–2016 outbreak [37], we estimated the proportion of reported cases (out of all known cases) in DRC to range from a low of 68% to a high of 96%, as extremes of the best known range of estimates, to be evaluated on their fit to data. Given these low and high estimates of reported cases and the estimate of vaccine effectiveness, a low estimate of vaccination program coverage was 44% (68% *×* 64.6%) and a high estimate of vaccination program coverage was 62% (96% *×* 64.6%). Vaccination was included in the simulation using a start date of August 14, 2018, estimated from dates available from situation reports [3].

To generate forecasts from each data set, 320 to 330 outbreaks were collected that passed the step of filtering on approximate agreement with DRC case counts. The simulated outbreaks that were retained after filtering were continued until they generated no further cases. Rare outlying simulated outbreaks that exceeded 300,000 cases were capped at the first value reached above that number, to limit unnecessary computation. We used this ensemble to derive a distribution of final outbreak sizes, and of cumulative counts at specific forecasting dates. Projection distributions were constructed using kernel density estimation with leave-one-out cross-validation to determine bandwidth, using a log-normal kernel for final sizes, due to the extended tail of the values, and a normal kernel for all other estimates. We calculated median values and 95% prediction intervals using the 2.5th and 97.5th percentiles of simulated outbreak size.

The frequencies of occurrence of the three vaccination coverage scenarios modeled (zero, low, and high coverage) among the simulated outbreaks accepted by the stochastic model’s filtering step were used to estimate likelihoods of the three scenarios. The frequencies of (*R*_initial_, *τ*) pairs selected by the filtering process were similarly recorded as an estimate of the likelihood of those transmission rate parameters.

We provide a detailed report of the parameters, simulations, and performance of the stochastic model in the Supplemental Material. This analysis was conducted using R (R Foundation for Statistical Computing, Vienna, Austria).

### Auto-Regression model

A negative binomial autoregressive model was chosen through a validation process to forecast additional new case counts at time points one week, two weeks, four weeks, and eight weeks from the current date. To adjust for disparities in the frequency of case reporting in historic outbreaks, the data were weighted by the inverse square root of the number of observations contributed to the model. Models considered included parameters for historic cumulative case counts (probable and confirmed) at different time points, logs of historic case counts, ratio of historic case counts to try and capture the trend of the epidemic curve, log(time), and an offset for current case total. When historic case counts for specific dates were missing, each missing case count was linearly interpolated from the two nearest case counts, allowing the model to remain agnostic about the current trend of the epidemic. After model fitting and validation, the final model chosen was a log-link regression for additional cases on the number of new cases identified in the previous two and four weeks and the ratio of these two case counts.

This analysis was conducted using R 3.4.2 (R Foundation for Statistical Computing, Vienna, Austria).

### Theil-Sen regression model

We conducted a simple regression forecast based solely on outbreaks of size 10 or greater, based on prior outbreaks [17–30]. Nonparametric Theil-Sen regression (R package mblm [38]) was used to project the final outbreak size based on values of the outbreak size at a specific earlier time. All time series were aligned on the day they reached 10 cases. Finally, we reported the median and 95% central coverage intervals for the prediction distribution, conditional on the predicted value being no smaller than the observed value for each day. More details can be found elsewhere [16].

This analysis was conducted using R 3.4.2 (R Foundation for Statistical Computing, Vienna, Austria).

### Gott’s rule model

Gott’s rule assumes we have no special knowledge of our position on the epidemic curve [39]. Given *Y*_0_ cases reported, assuming a non-informative uniform prior for the portion *α* of the epidemic observed to date, the corresponding probability density function for the final size Y = *Y*_0_*/α* is *Y*_0_*/y*^2^, *Y*_0_ ≤ *y*. We constructed a probability mass function by assigning all probability density to the whole number of days given by the integer part of each value. We used this probability mass function as a projection of the final outbreak size.

This analysis was conducted using R 3.4.2 (R Foundation for Statistical Computing, Vienna, Austria).

### Scoring

Models’ performance on short-term projections was scored using the natural logarithm of the probability assigned to subsequently known reported case counts.

The stochastic and negative binomial auto-regression models were scored based on projections at multiple time points, where available:

- Case count 1 week after the last available case count
- Case count 2 weeks after the last available case count
- Case count 4 weeks after the last available case count
- Case count 8 weeks after the last available case count

For each projection, we generated an assignment of probability to possible values of multiple quantities. As Ebola situation reports were released before February 25, we generated short-term projections in real time with stochastic and negative binomial auto-regression models. We scored these projections and calibrated the models during the outbreak. Then we used these calibrated models for our projections on February 25. The Theil-Sen and Gott’s rule models were not calibrated as the outbreak’s final size is not known. Final outbreak size projections generated by the stochastic, Theil-Sen, and Gott’s rule models were recorded for future evaluation of their performance.

### Evaluation of a catastrophic outbreak

Because the question has been raised of whether the current outbreak might grow to the scale of the catastrophic West Africa outbreak, we evaluated each model’s projected probability of exceeding three large outbreak scenarios of 1,000 cases, 10,000 cases, or the 28,616 cases reported in the West Africa outbreak [30]. Each of these three thresholds is more than double the size of the previous second largest outbreak (*n*=425, Uganda, 2000) [22].

## Results

When we started performing our short-term forecasts on August 20, 2018, there were 102 reported EVD cases in North Kivu and Ituri provinces, DRC. We used our stochastic and auto-regression models to project one-week, two-week, four-week, and eight-week forecasts of outbreak size. As time lapsed, we compared predicted and known outbreak sizes and found a higher probability of accurate forecasts at one week than at eight weeks (Figs 2, 3). Log-likelihood scores typically declined as projection times extended further into the future. These lower-scoring longer-term projections tended to include wider prediction intervals, reflecting less certain outcomes in which less probability was assigned to any one value. The epidemic curve accelerated in early October, and stochastic model forecasts occurring just before that change were especially low scoring, as they failed to anticipate the coming rise in case counts. Subsequent projections took into account the reported acceleration and their performance recovered.

**Fig 2.**
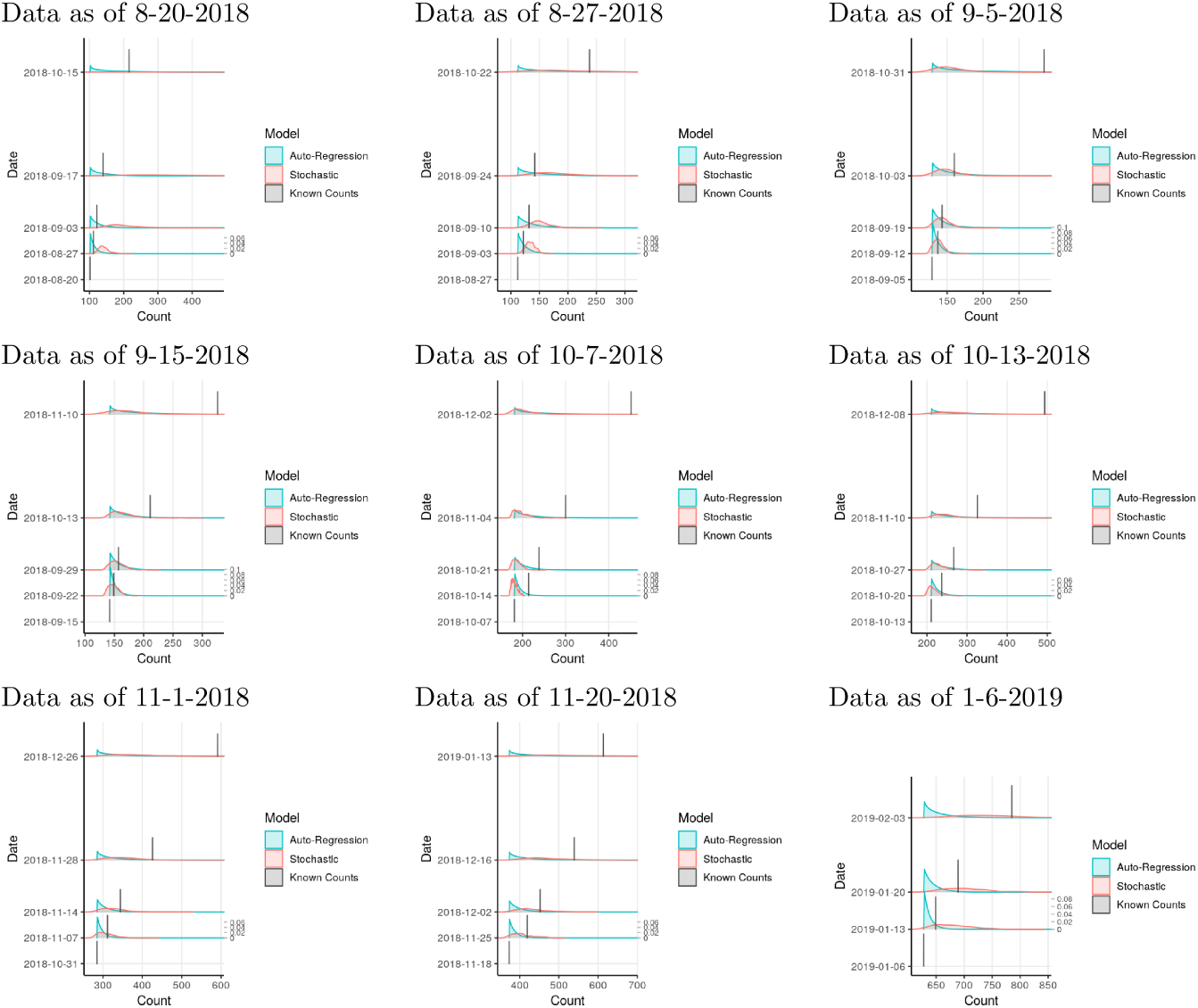
**Prospective validation of probabilistic projections** of auto-regression and stochastic models, by comparing projected cumulative case counts from past data to known case counts. Vertical lines indicate known case counts (height not to scale).

**Fig 3.**
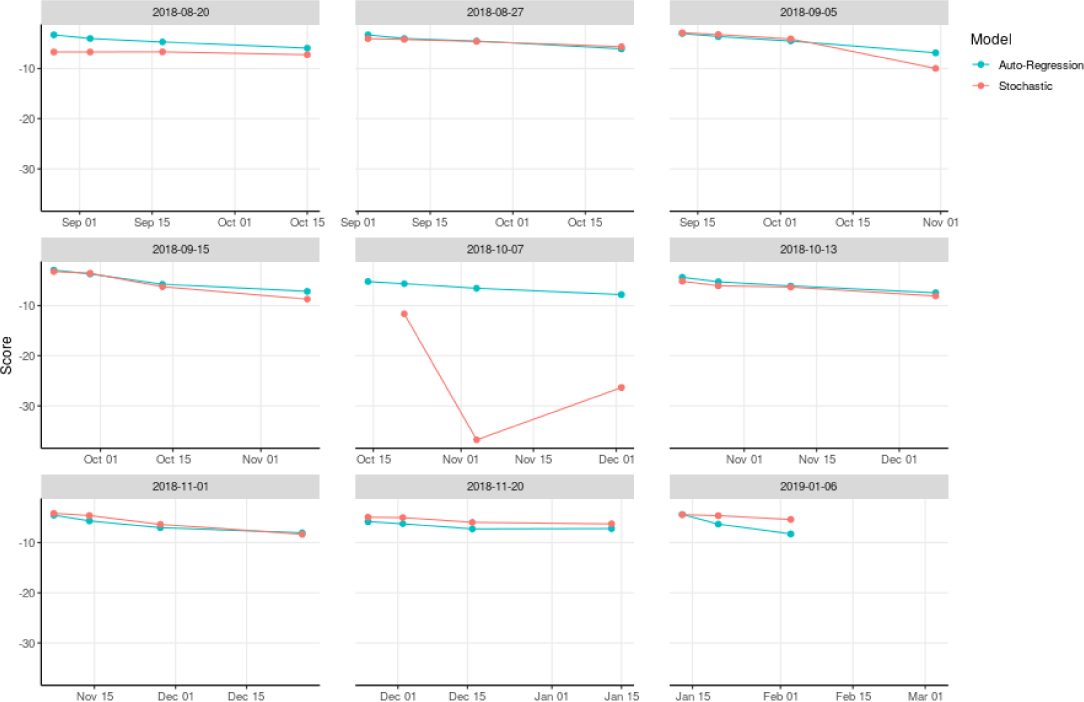
Log-likelihood scores of stochastic and auto-regression projections in prospective validation.

After our model validation process was completed, we used the stochastic and auto-regression models to project one-week, two-week, four-week, and eight-week outcomes (Figs 4, 5). We used the Gott’s rule and Theil-Sen regression models together with the stochastic model to project final outbreak sizes (Figs 6, 7).

**Fig 4.**
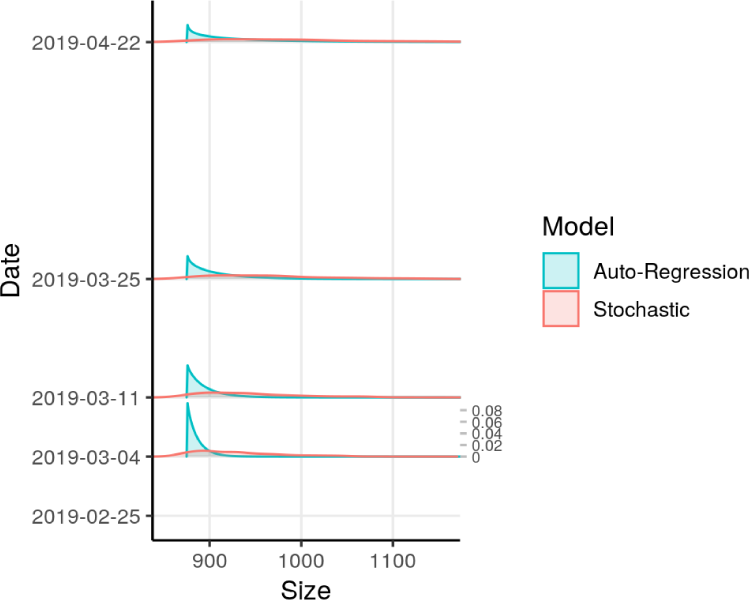
**Short-term probabilistic projections of case counts** based on reported counts as of February 25, 2019. Stochastic and auto-regression models were used. Vertical scale of probabilities, on right side of plot, applies to all dates.

**Fig 5.**
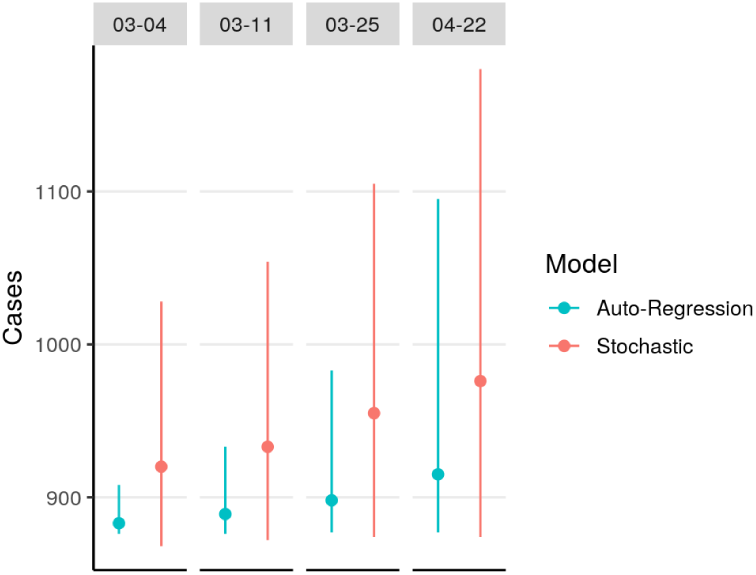
**Medians and prediction intervals from short-term projections of case counts** based on reported counts as of February 25, 2019. Solid dots indicate median count.

**Fig 6.**
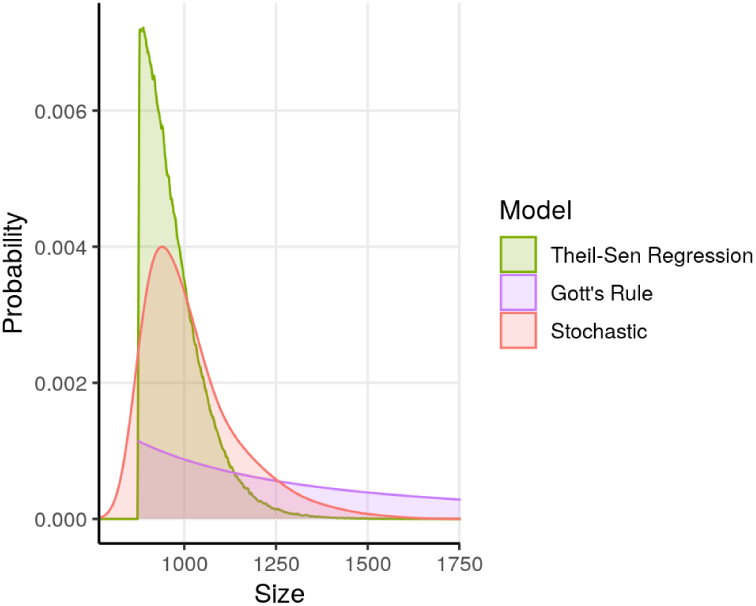
**Probabilistic projections of final case counts** based on reported counts as of February 25, 2019. Stochastic, Theil-Sen regression, and Gott’s rule models were used.

**Fig 7.**
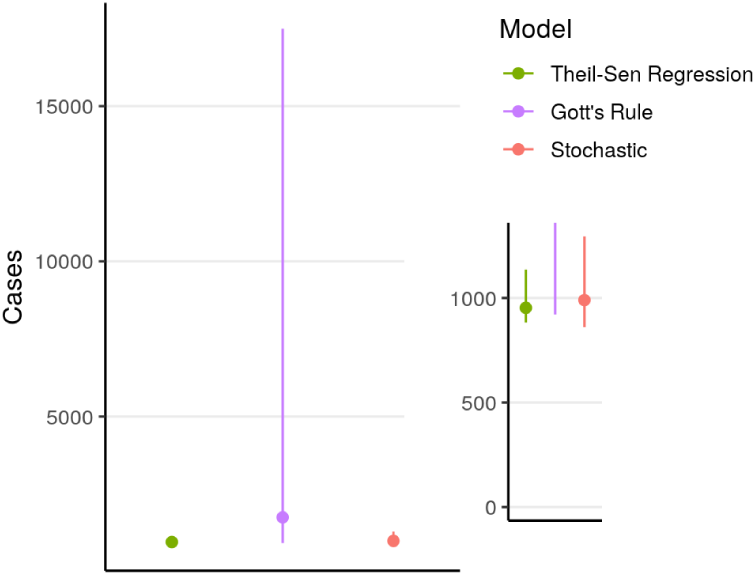
**Medians and prediction intervals from projections of final case counts** based on reported counts as of February 25, 2019. Solid dots indicate median size. Inset provides detail at smaller numbers.

As of February 25, the total reported count of probable and confirmed EVD cases was 875 cases. With the stochastic model, the four-week projection of median outbreak size was 955 cases (95% prediction interval: 874–1105). With the auto-regression model, the four-week projection of median outbreak size was 898 cases (95% prediction interval: 877–983).

Median final outbreak size projected by the stochastic model was 990 cases (95% prediction interval: 860–1295). With the Gott’s rule model, the median final outbreak size was 1749 cases (95% prediction interval: 921–17498). Median final outbreak size projected by the regression model was 953 cases (95% prediction interval: 882–1136).

The possibility of a final outbreak size exceeding 1,000 cases was projected to be quite likely by all the models, with the Gott’s rule forecast being particularly pessimistic, as it was in all cases. The probability of a very large outbreak (10,000 or more cases) was calculated to be below 8% for Gott’s rule and negligibly small for the other models. A final outbreak size of a catastrophic outbreak larger than the West Africa outbreak (28,616 or more cases) was projected to have probability less than 3% by Gott’s rule and negligibly small for the other models (Table S2 in S1 Supporting Information).

We generated short- and long-term projections from earlier snapshots of the current outbreak case counts with each model, for the purpose of both scoring and forecasting. These additional results can be found in the Supplemental Material.

### Probability of vaccination coverage scenarios

The stochastic model estimated likelihoods of the three scenarios of zero, low, and high vaccination coverage, based on how often models using the different coverage assumptions were able to pass the particle filtering step. In the estimate based on data through February 25, vaccination coverage was estimated substantially more likely than low or high coverage, as was also true in most earlier estimates. The lower vaccination coverage scenario was estimated more likely than the higher one. Higher vaccination coverage scenarios were estimated more likely in estimates made before October, at which time the outbreak epicenter shifted into the conflict zone and situation reports described an increase in case counts (Fig S5 in S1 Supporting Information). Assuming that other transmission factors were held constant, our result may provide evidence that that geographical shift in transmission contributed to a decreased likelihood of a high vaccination scenario.

## Discussion

As of February 25, 2019, with 875 cases reported to date, most of our model projections were concentrated in a range up to about 300 additional cases overall, and even the Gott’s rule forecast, whose prediction intervals are especially wide, placed low probability on outcomes on the scale of the West African outbreak. The current outbreak, however, has so far behaved unpredictably, possibly due to complex social circumstances, and could continue to be unpredictable and render our projections inaccurate. EVD has never before been introduced into a conflict zone with such political instability, potential disease mobility and community impenetrability [2]. In October, WHO reported that up to 80% of contacts were not being traced [3]. At present, the most reliable data source of EBOV transmission has been the weekly case counts that can be found in the WHO situation reports. Despite such situations of data scarcity and new outbreak circumstances, our models generated relatively accurate short-term projections of outbreak size in the months of January and February 2019, suggesting that short-term projections made in real time can be useful in decision making and resource allocation.

Another outbreak response challenge has been ascertaining the level of vaccination coverage that has occurred during the ring vaccination program. For the ring vaccination program to be theoretically effective at reducing transmission, an uncertain proportion of contacts and contacts of contacts need to receive the vaccine [6]. We estimated that transmission rates are higher than would be expected under target levels of 62% coverage due to contact tracing and vaccination. Under our stochastic model’s assumptions, the elevated transmission rates, relative to those of past outbreaks, needed to reproduce this outbreak’s epidemic trajectory are highly unlikely to be randomly observed with high vaccination coverage. This model estimate may serve as a loose proxy for the outbreak response challenges. Thus, outbreak response teams may need to consider other control and care strategies to end the outbreak. Under current circumstances, mass vaccination in regions of high prevalence or areas where the outbreak has newly arrived may be a more effective strategy to reduce EBOV transmission than ring vaccination. Aggressive supportive care, experimental therapeutics and high-quality facilities (*e.g.* air-conditioned, individualized) have also been described as part of the outbreak response. These interventions have the potential to improve health-seeking behaviors and possibly reduce EBOV transmission in communities that are resistant to control efforts [15], but further studies are needed.

The mathematical models we adopted for this project near the beginning of the outbreak were developed to predict the course of prior outbreaks, and did not consider circumstances unique to the current outbreak. To improve the accuracy of short-term projections, we used the short-term forecasting performance of the mathematical models during the current outbreak to calibrate them and improve their performance on this outbreak. This may be a useful pattern for short-term forecasting of ongoing disease outbreaks in real time.

The performance of our short-term prediction models can be assessed on the outbreak to date. The relatively simple auto-regression model we used performed more consistently on the range of partial data sets used for scoring than the more complex stochastic simulation model did. The stochastic model has tended to produce tighter prediction distributions that are prone to extreme failure when they get it wrong, while the auto-regression model’s predictions are more tolerant of unpredictable outcomes. Conversely, the stochastic model outperforms the auto-regression when it gets the prediction right. It should also be noted that because the stochastic simulation model is based on mechanistic knowledge of the transmission process generating the outbreak while the regression model is a purely statistical inference from past outbreaks, it may be that if conditions emerge that are substantially different from past outbreak conditions, the mechanistic model may produce sensible predictions where a purely statistical model fails. Because the other models included produce only final outbreak size projections, they can not be fully evaluated before the outbreak has ended. However, we can note that our early projections of final size (Figs S9, S10 in S1 Supporting Information) fell below the counts that have been observed as of February. In other words, our forecasts based on early reports and on an assumption that past outbreaks can be used to forecast the present one were more optimistic than warranted by subsequent events. Events to come may shed light on whether this outbreak is qualitatively different than the past ones we have used to construct forecasts, perhaps due to the impact of conflict conditions on the outbreak.

There are limitations to our projections. Projection distributions are right-skewed, with long tails (and we therefore report the median instead of the mean). We were unable to include all the 23 observed EVD outbreaks with a case count greater than ten cases in our estimates due to data availability. Our regression models are based entirely on past outbreaks of Ebola virus disease (measured and reported in different ways), and cannot account for the improved control measures and vaccination in the way that a mechanistic model does. We included as much real-time information in our models as possible, but situations such as the introduction of EVD into a zone of armed conflict and the recent introduction of vaccination are not reflected by the suite of past outbreaks. The stochastic model used estimated vaccination effectiveness, reported cases, and timing of onset dates affected by vaccination from studies from West Africa, not DRC, and did not include vaccination of healthcare workers. Our forecasts do not account for possible unreported cases or changes in reporting over time; such gaps in reporting can not be ruled out, though given the intense efforts at case finding that started in July and carried forward, we think it unlikely that there were large changes in reporting. Furthermore, as the outbreak moved into areas affected by violent conflict as the outbreak continued, we think it likely that case reporting, if anything, decreased over time, and therefore underreporting would not explain the apparent increase in transmission from June to October. While it would be desirable to use the vaccination coverage estimates to estimate the number of cases prevented by the vaccination program, our models were not designed to produce a testable estimate of such an effect.

Gott’s rule [39], presented initially as reflecting a “Copernican” principle with respect to time, assumed that ones observation of the age of a phenomenon occurred at a random time during its lifetime. Gott applied this principle in a variety of settings, including the lifetime of Broadway shows, the Berlin Wall, and of the human race [39, 40]. Critics suggest that Gott’s rule gives misleading predictions when the phenomenon in question exhibits known time scales, such as in prediction of human lifetimes [40, 41]. Gott’s rule has not been validated for epidemics, and in this case, is based on an assumption that the current case count in an epidemic occurs at a random point in its progression. For an easily detected but uncontrollable supercritical epidemic, we would expect forecasts based on the first reports to underestimate the final size. For an easily controlled epidemic, forecasts would more likely be too pessimistic, since control measures would (by assumption) swiftly end the epidemic after recognition. Our application of the rule is intended to serve as a null hypothesis-like comparator for the other methods.

Our models do not account for spatial heterogeneity in transmission, which may be relevant to the course of this outbreak, in which an increase in transmission appears to coincide with a shift in location. The stochastic model may be sensitive to the assumption of an exponentially decaying curve for transmission rate by day, and alternative assumptions might lead to a different distribution of forecast outcomes. Indeed, all of our projections are conditional on model assumptions being met. If unpredictable events were to change patterns of EBOV transmission in ways not seen in past outbreaks, assumptions used for model projections could be violated and our results could change. For example, exceedingly improbable events such as a catastrophic outbreak (more than 10,000 cases or approaching the size of the 2013–2016 West Africa outbreak) might become more probable. A strength of our approach was the use of multiple methods to estimate the outbreak size, including both mechanistic modeling and purely statistical approaches.

Before October, there was limited EBOV transmission in active, armed conflict zones. When more reported EVD cases occurred in Beni, it was unclear how the context would affect secondary EBOV transmission. The October data suggests that EBOV transmission increased there. We believe the increased rate of reported EVD cases corresponded to the shift of EBOV transmission into conflict zones. It may be that a model that explicitly distinguishes transmission rates in these zones from those in other areas would model the dynamics underlying these cases more precisely and produce more accurate projections.

Many of the reported EVD cases in this outbreak have been identified as new transmission chains. While a reduced ability to detect and vaccinate contacts in difficult-to-access communities has been anecdotally reported, our stochastic model provided empirical evidence of outbreak challenges, suggesting that transmission rates were consistent with lower than target levels of vaccine coverage. beyond those already reported. While a catastrophic outbreak of 10,000 or more cases is not projected as probable, vigilance is warranted. New circumstances—such as epidemic spread to Uganda—call for newly validated projections, whenever possible, even in the short term.

## S1 Supporting Information. Appendix containing supplemental figures and tables

### Supporting information

Table S1 summarizes the past outbreaks used as data to inform our models.

**Table S1.** Table of past outbreaks by year and country.

| Time Period | Country | Reported Count | Time Series Count | Regression? | Stochastic? | Auto-Regression? |
| --- | --- | --- | --- | --- | --- | --- |
| Aug–Sep 1976 | DRC* | 318 | 262 | Yes | No | Yes |
| Jun–Nov 1976 | Sudan | 284 | 284 | Yes | No | Yes |
| Aug–Sep 1979 | Sudan | 34 | 34 | Yes | Yes | Yes |
| Dec 1994–Feb 1995 | Gabon | 52 | 49 | Yes | No | Yes |
| May–Jul 1995 | DRC | 315 | 317 | Yes | Yes | Yes |
| Jan–Apr 1996 | Gabon | 37 | 29 | Yes | Yes | Yes |
| Jul 1996–Mar 1997 | Gabon | 60 | – | No | No | No |
| Oct 2000–Jan 2001 | Uganda | 425 | 436 | Yes | No | Yes |
| Oct 2001–Jul 2002 | Gabon, Republic of the Congo | 124 | 124 | Yes | Yes | Yes |
| Dec 2002–Mar 2003 | Republic of the Congo | 143 | – | No | No | No |
| Nov–Dec 2003 | Republic of the Congo | 35 | 35 | Yes | Yes | Yes |
| Apr–Jun 2004 | Sudan | 17 | 17 | Yes | Yes | Yes |
| Apr–May 2005 | DRC | 12 | 12 | Yes | Yes | Yes |
| Aug–Nov 2007 | DRC | 264 | 264 | Yes | Yes | Yes |
| Dec 2007–Jan 2008 | Uganda | 131 | 127 | Yes | Yes | Yes |
| Dec 2008–Feb 2009 | DRC | 32 | 32 | Yes | Yes | Yes |
| Jun–Aug 2012 | Uganda | 24 | 24 | Yes | Yes | Yes |
| Jun–Nov 2012 | DRC | 52 | 52 | Yes | Yes | Yes |
| Aug–Nov 2014 | DRC | 66 | 62 | Yes | Yes | Yes |
| Jul–Oct 2014 | Nigeria (offshoot of West African outbreak) | 20 | – | No | No | No |
| Jan 2014–Jun 2016 | Guinea, Liberia, Sierra Leone | 28,616 | 21,422 | Yes | No | Yes |
| Apr–Jun 2018 | DRC | 53 | 53 | Yes | Yes | Yes |

Official reported case counts for each epidemic are given, including suspected cases (“Reported Count”). Case counts for the time series data included in the models include only probable and confirmed cases (“Time Series Count”). Case counts for historic outbreaks were pulled from publicly available literature [17–30]. Lastly, each historic outbreak’s inclusion in the regression, stochastic, and auto-regression models is enumerated.
\*Democratic Republic of Congo (formerly Zaïre)

We retained snapshots of the set of available case counts at multiple time points, for use in scoring of retrospective model projections against known subsequent counts (Fig S1). In later data sets, due to the larger number of data points, a subset of the case counts was selected for use in the stochastic model’s particle filtering step, as noted in the figure.

#### Stochastic Model

**Fig S1.**
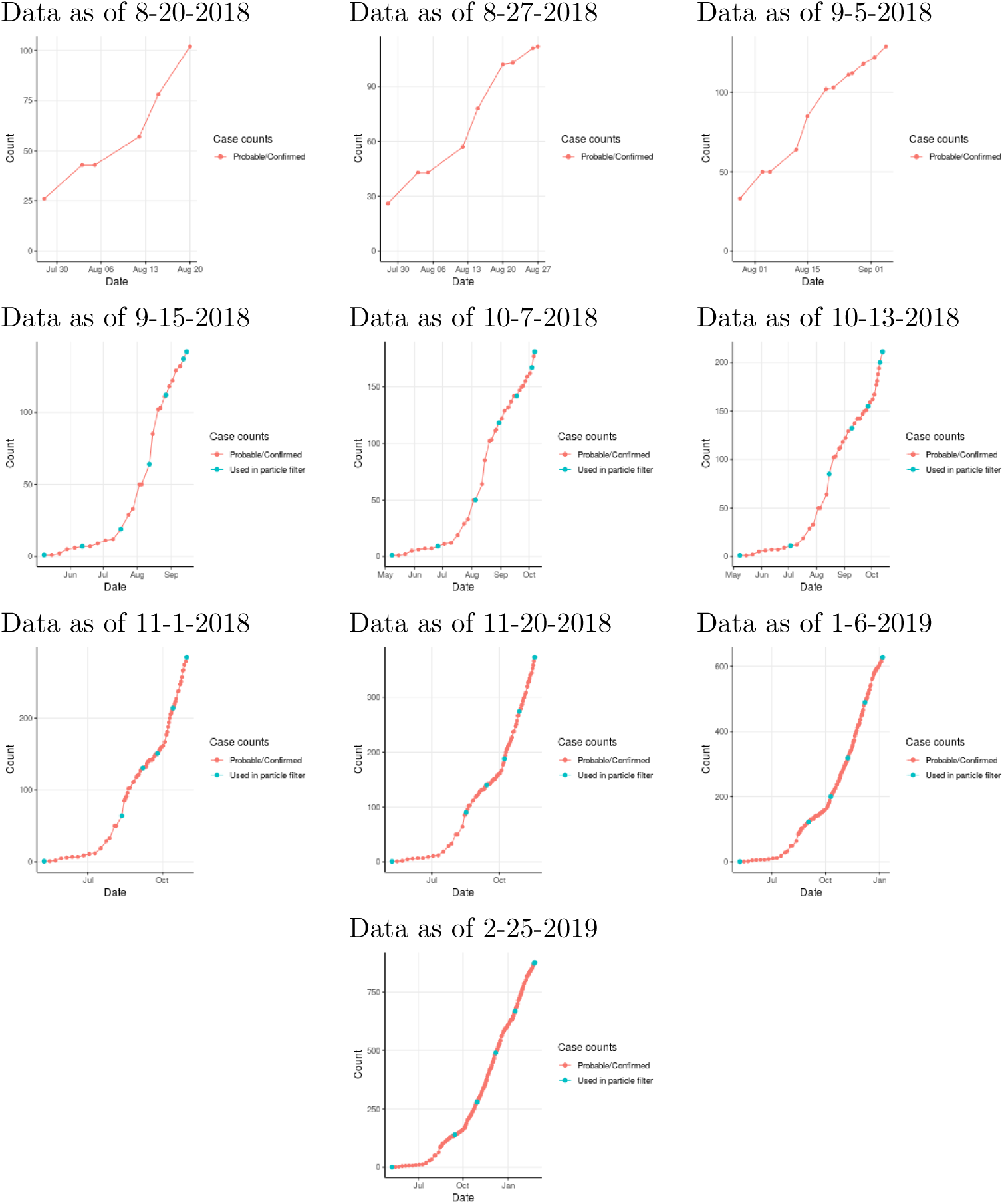
**Reported case counts in current outbreak** by date, in multiple snapshots of available data. Where not otherwise noted, all case counts shown were used in the stochastic model’s particle filtering step.

Epidemic curves reported for past Ebola outbreaks were used to estimate time series of effective reproduction number (*R*) by day, which were then fit to an exponential quenching curve (Fig S2). The quenching parameter estimates the rate of change in *R* per day from *R*_initial_ that results from outbreak control efforts, behavioral changes in response to the outbreak, and potential local depletion of susceptibles. The parameters *R*_initial_ and *τ* estimated by that curve fitting on past epidemics were then used to create a distribution from which values were sampled to parametrize the stochastic simulation (Fig S3).

**Fig S2.**
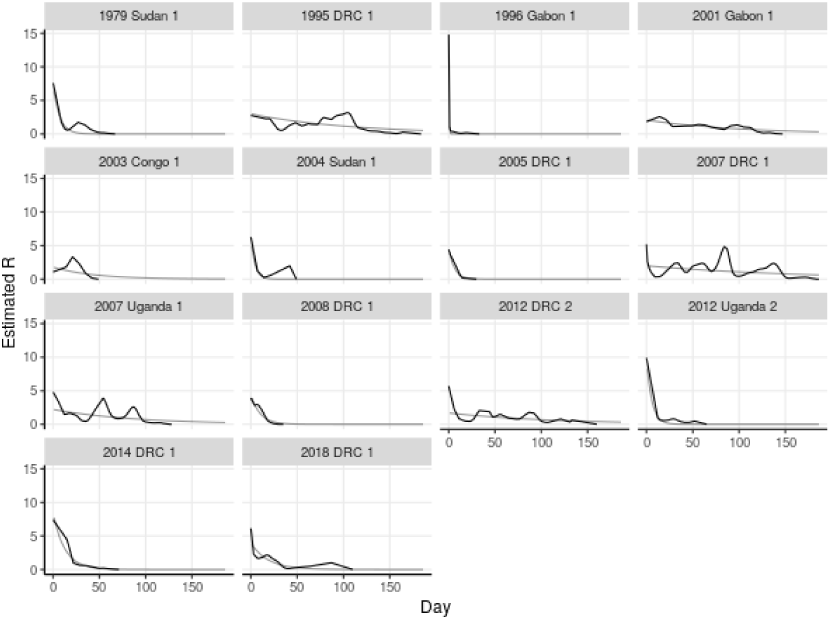
Estimates of reproduction number. *R* by day in past Ebola outbreaks. Thin curves are exponentially quenched curves *R* = *R*_initial_*e*^*-τd*^ fit to each series of *R* estimates.

**Fig S3.**
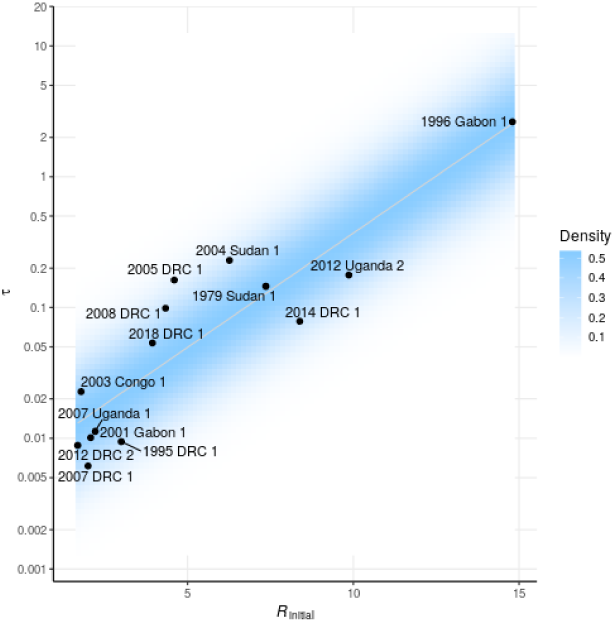
Distribution of transmission rates sampled for simulation. Black dots are pairs *R*_initial_ and quenching rate *τ* estimated from past Ebola outbreaks, and blue cloud is the continuous distribution from which pairs are sampled for simulation.

The *R*_initial_ and *τ* parameters driving simulated outbreaks that were successful in passing the particle filtering step tended to cluster in particular locations within the assumed distribution (Fig S4). In some cases, distinct ranges of *R*_initial_ and/or *τ* were selected in conjunction with the different vaccine coverage scenarios. The frequencies of selection of the three vaccine coverage scenarios were used to estimate the likelihood of each scenario (Fig S5).

For each data set, simulations were generated until between 320 and 330 simulated outbreaks were obtained after filtering based on approximate agreement with reported case counts from the current outbreak.

The simulations passing the particle filtering step, representing a distribution of parameter values and vaccine scenarios, were continued beyond the particle filtering points to generate a spreading set of projections of case counts at later dates (Fig S6), which was smoothed to create probabilistic projections of future case counts at the desired dates.

#### Projections

We have recorded the projections generated by our models from older data sets to assess the development of the projections as the outbreak has progressed, in Figs S7, S8, S9, and S10.

Table S2 summarizes the medians and 95% prediction intervals produced by each model on the most recent data set included, and their probabilities of large outcomes.

**Table S2.** **Table of medians and prediction intervals** of model projections.

| Forecast as of | Forecast | Model | Lower | Median | Upper | Size 1,000 | Size 10,000 | Size 28,616 |
| --- | --- | --- | --- | --- | --- | --- | --- | --- |
| 2019-02-25 | 2019-03-04 | Auto-Regression | 876 | 883 | 908 | 0.00002 | 0.00000 | 0.00000 |
| 2019-02-25 | 2019-03-04 | Stochastic | 868 | 920 | 1028 | 0.11077 | 0.00000 | 0.00000 |
| 2019-02-25 | 2019-03-11 | Auto-Regression | 876 | 889 | 933 | 0.00204 | 0.00000 | 0.00000 |
| 2019-02-25 | 2019-03-11 | Stochastic | 872 | 933 | 1054 | 0.16451 | 0.00000 | 0.00000 |
| 2019-02-25 | 2019-03-25 | Auto-Regression | 877 | 898 | 983 | 0.03327 | 0.00000 | 0.00000 |
| 2019-02-25 | 2019-03-25 | Stochastic | 874 | 955 | 1105 | 0.26913 | 0.00000 | 0.00000 |
| 2019-02-25 | 2019-04-22 | Auto-Regression | 877 | 915 | 1095 | 0.16126 | 0.00000 | 0.00000 |
| 2019-02-25 | 2019-04-22 | Stochastic | 874 | 976 | 1180 | 0.39925 | 0.00000 | 0.00000 |
| 2019-02-25 | final | Gott’s Rule | 921 | 1749 | 17498 | 0.86625 | 0.07875 | 0.02183 |
| 2019-02-25 | final | Stochastic | 860 | 990 | 1295 | 0.46620 | 0.00000 | 0.00000 |
| 2019-02-25 | final | Theil-Sen Regression | 882 | 953 | 1136 | 0.29805 | 0.00000 | 0.00000 |

**Fig S4.**
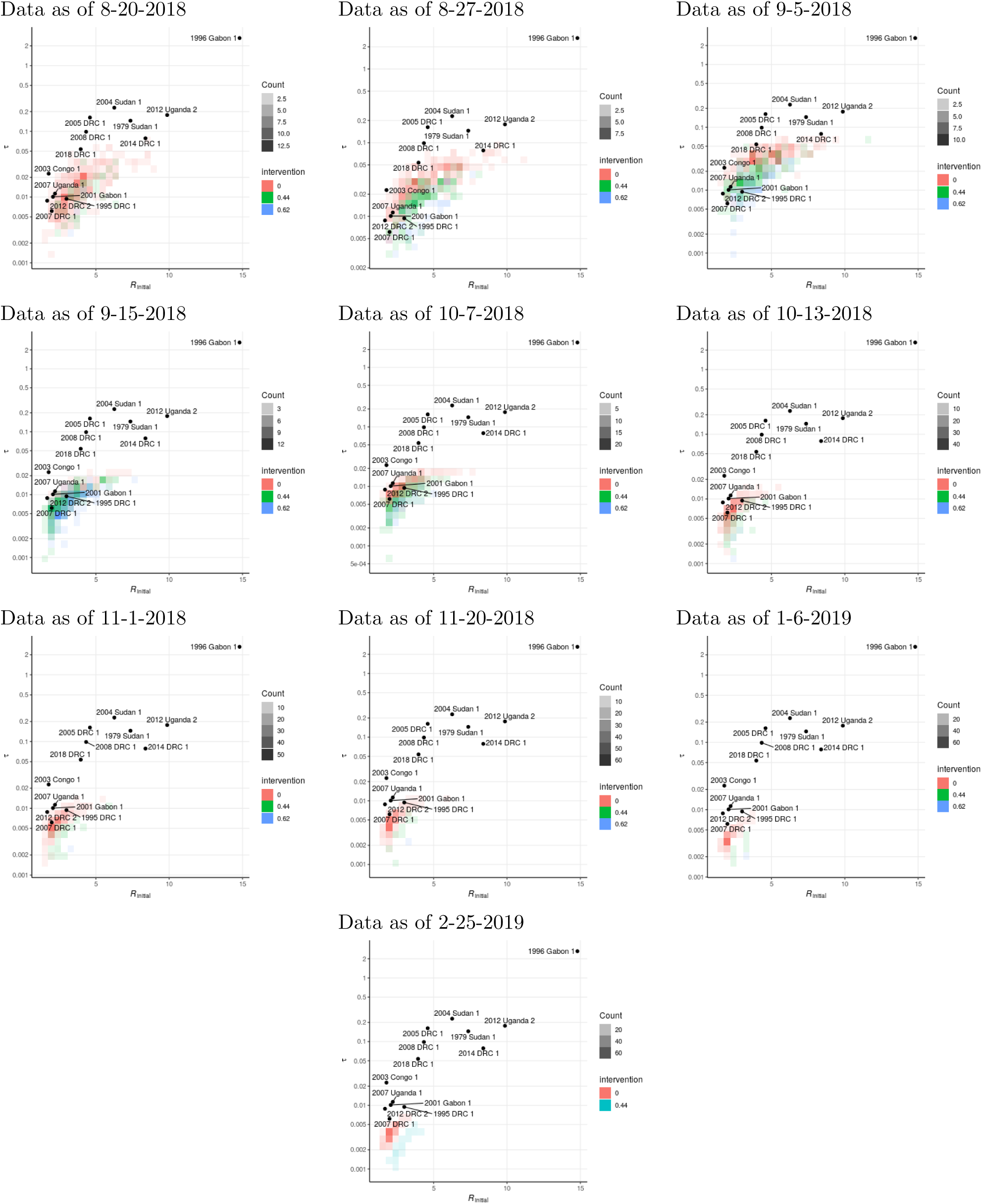
**Transmission rates selected by the particle filtering process,** by vaccine coverage scenario, for successive snapshots of available case count data. As in previous figure, black dots for *R*_initial_, *τ* pairs estimated for past outbreaks (for comparison), and colors illustrate the density of *R*_initial_, *τ* pairs selected by filtering simulated outbreaks, classified by level of vaccine coverage.

**Fig S5.**
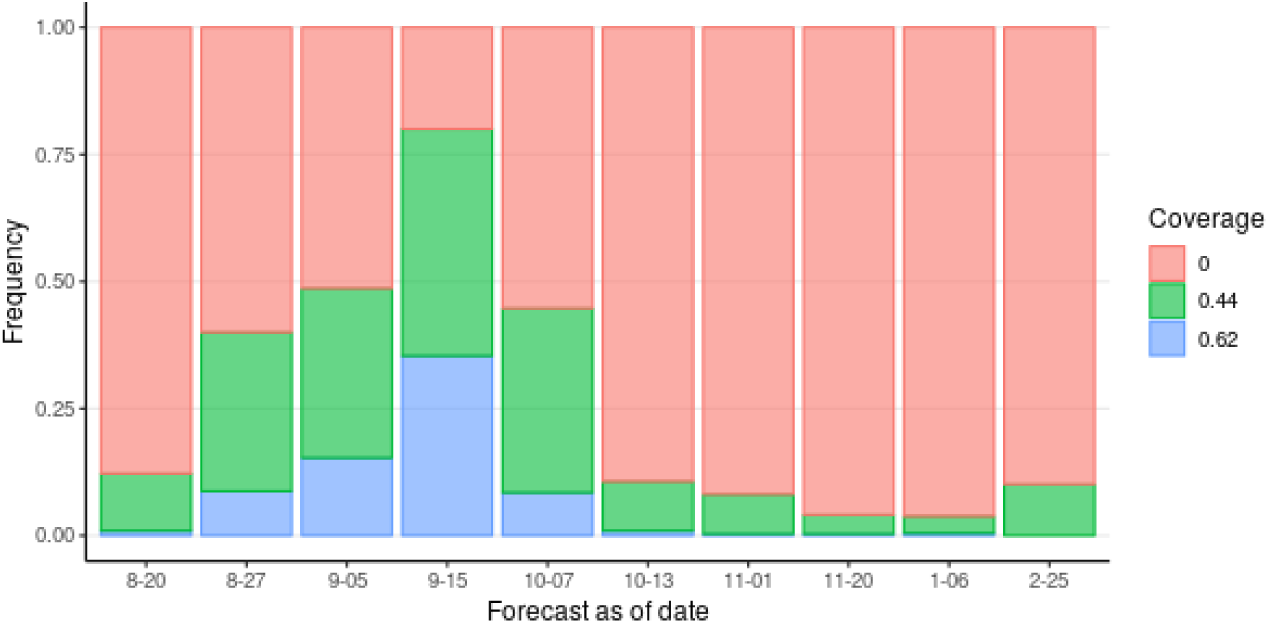
**Likelihood estimates of vaccine coverage scenarios consistent with model transmission rates,** estimated by number of simulated outbreaks under the three scenarios accepted by the stochastic model’s filtering step, in which simulated outbreaks must match reported case counts. Note that each bar of this plot is an estimate based on the entire time series up to the date indicated, that is, they are cumulative estimates of the overall vaccine coverage, not successive estimates of short-term coverage.

**Fig S6.**
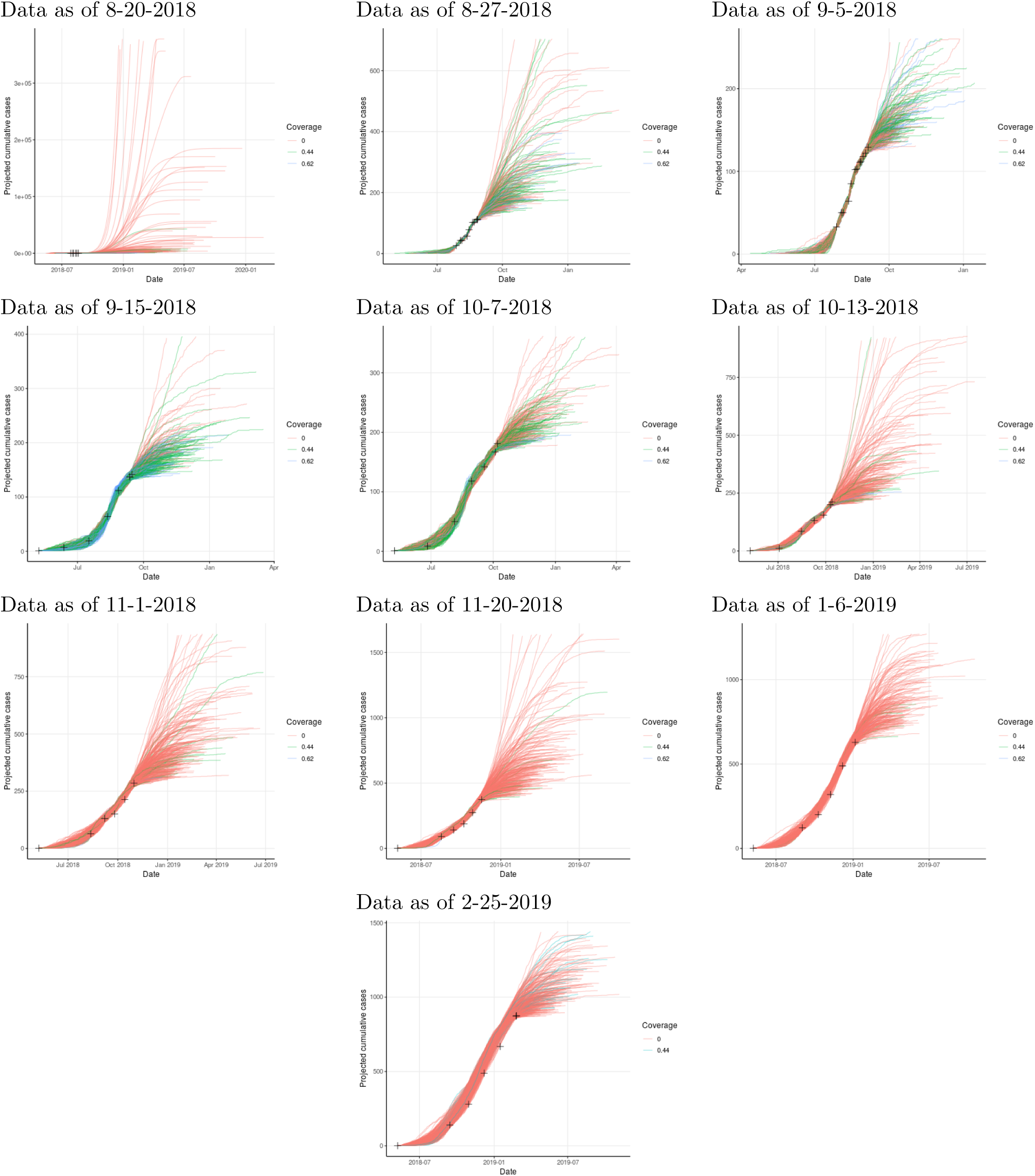
**Cumulative case counts by date** projected by individual realizations of the stochastic model, by vaccine coverage scenario, using successive snapshots of available case count data. The vertical axis is cut off at the upper limit of the 95% prediction interval for outbreak sizes, for readability.

**Fig S7.**
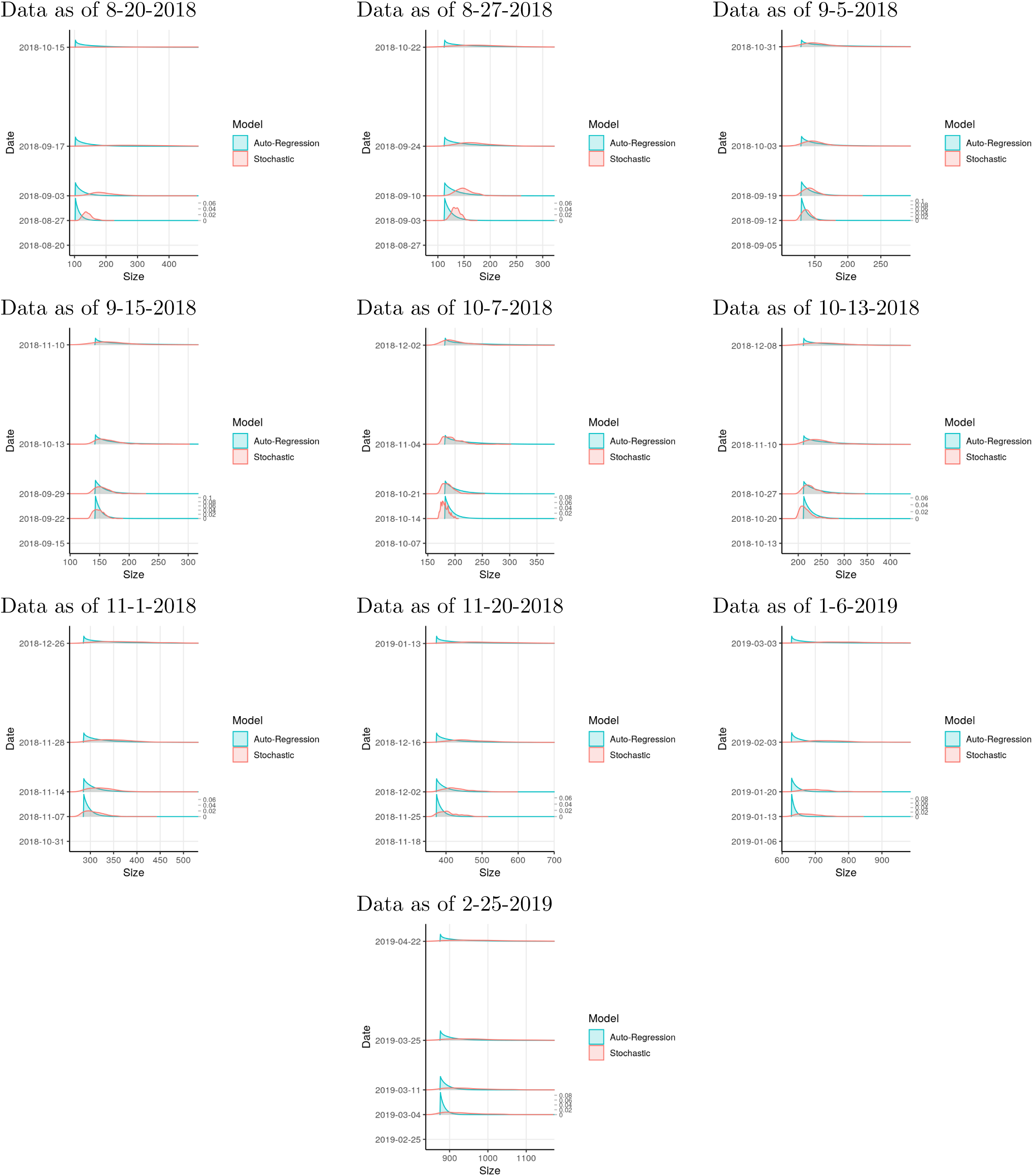
**Short-term projections** based on multiple data sets. Probability scale (marked on right) is consistent across dates within each subplot.

**Fig S8.**
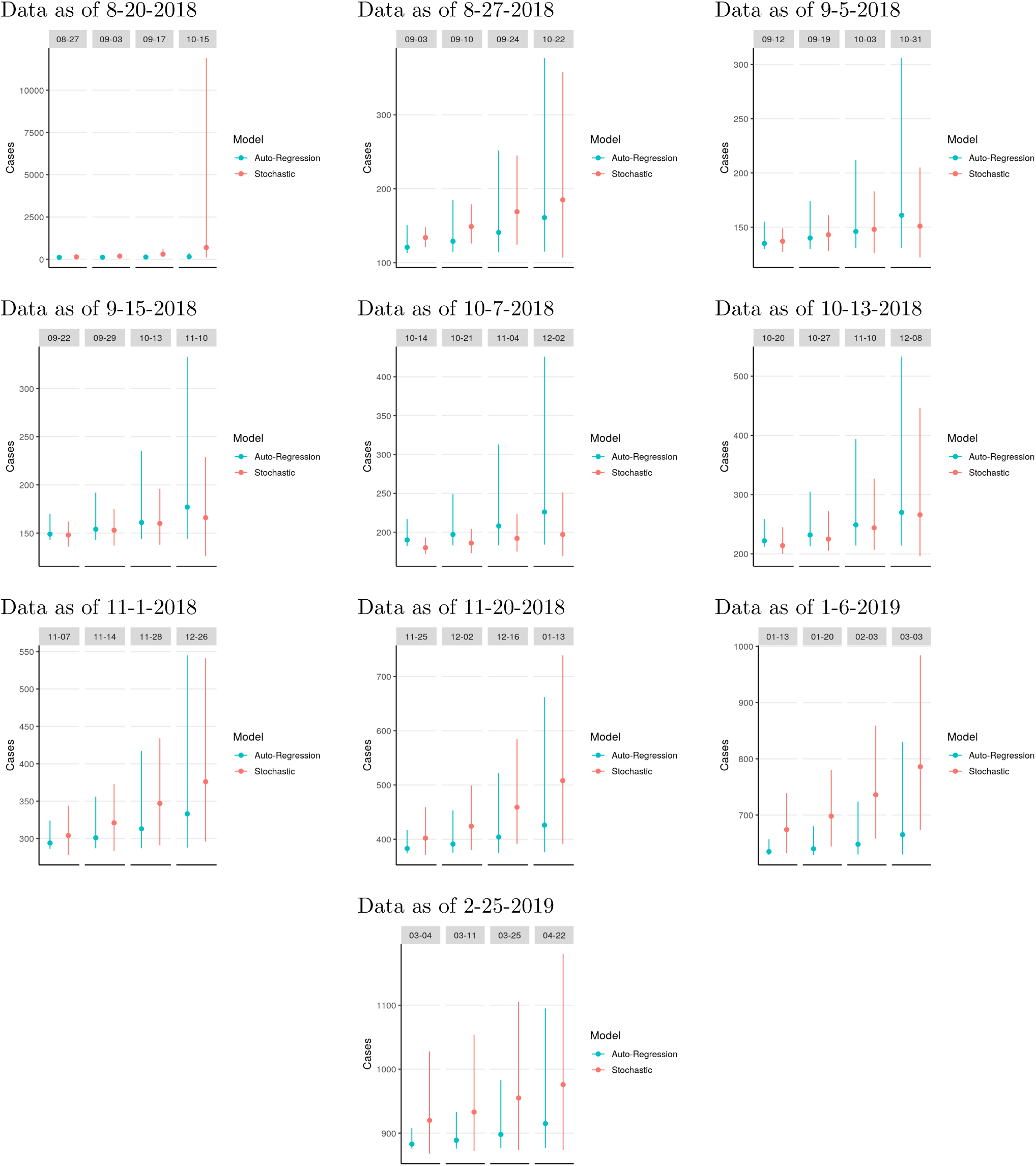
**Medians and prediction intervals from short-term projections** based on multiple data sets.

**Fig S9.**
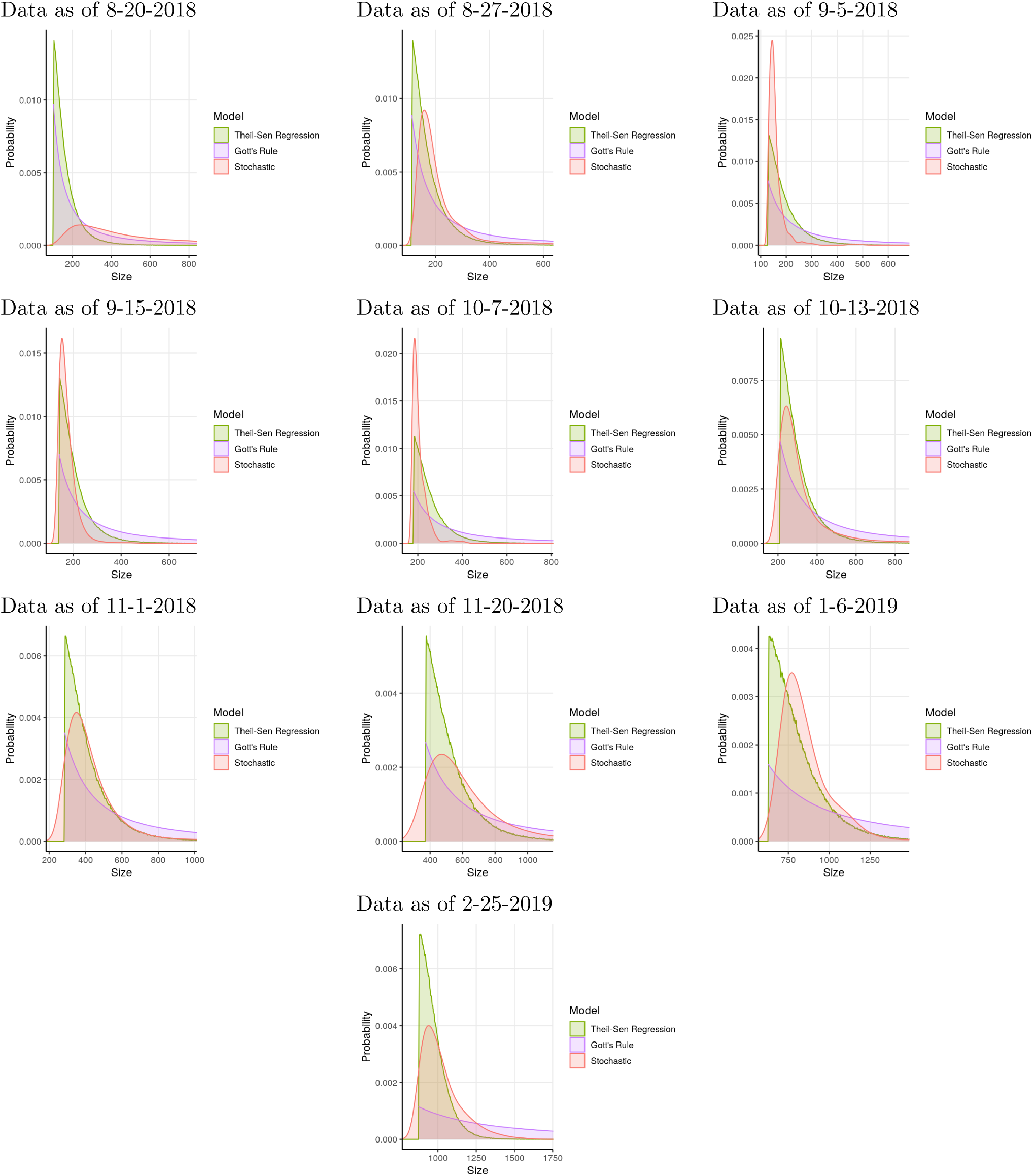
**Final outbreak size projections** based on multiple data sets.

**Fig S10.**
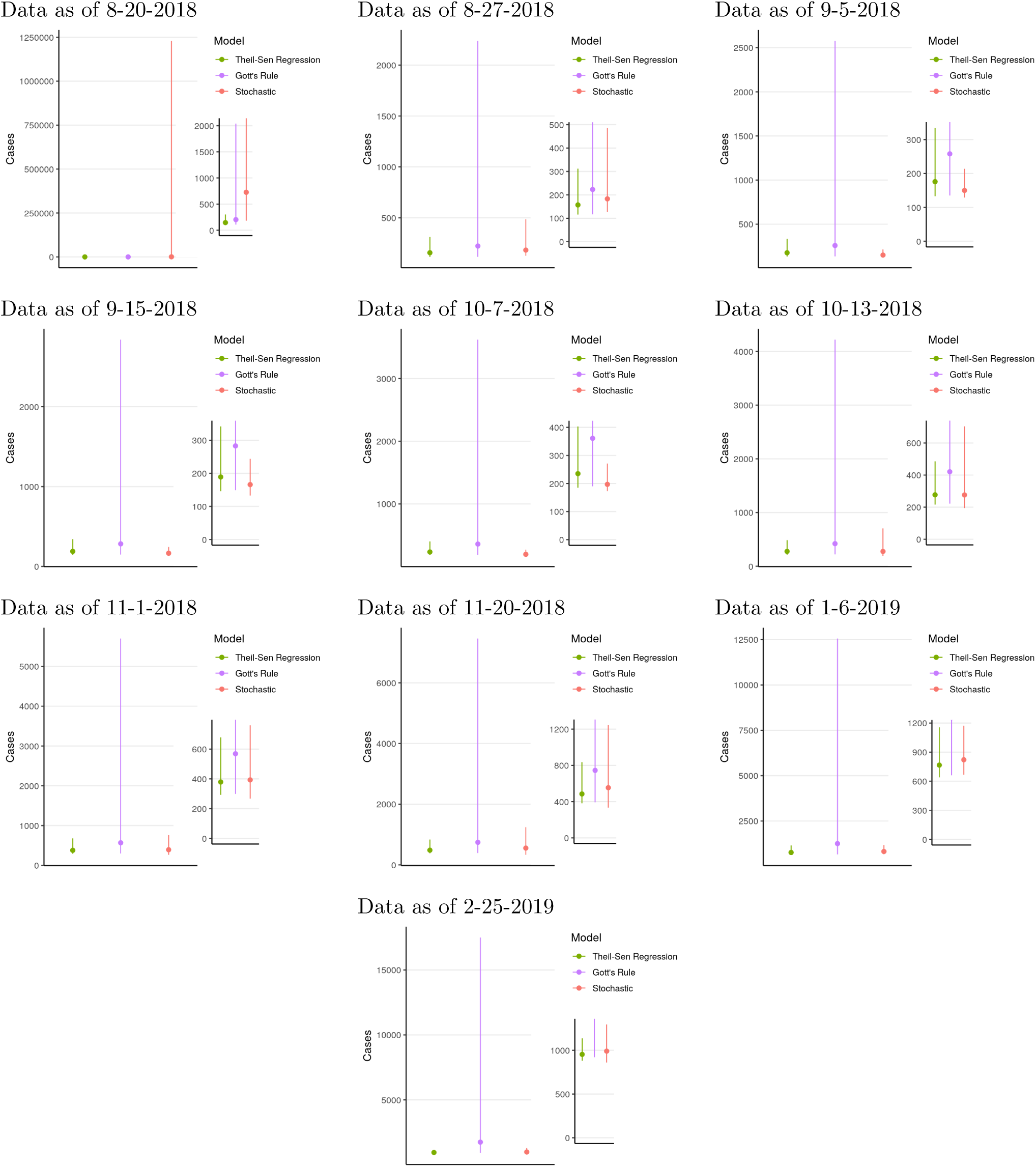
**Medians and prediction intervals from final outbreak size projections** based on multiple data sets, with inset for detail in smaller numbers.

